# Ventral Pallidum GABA Neuron Inhibition Augments Context-Appropriate Defensive Responses to Learned Threat Cues

**DOI:** 10.1101/2025.08.28.672918

**Authors:** Erica M. Ramirez, Maricela X. Martinez, Ryan K. Rokerya, Vanessa Alizo Vera, Christina M. Ruiz, Mitchell R. Farrell, Shreeya A. Walawalkar, Hazael Ramirez-Ramirez, Grace J. Kollman, Stephen V. Mahler

## Abstract

The ventral pallidum (VP) is embedded within the brain circuits controlling motivated behavior, which are heavily implicated in addiction and other psychiatric disorders. Prior work showed that VP GABAergic neurons (VP^GABA^) promote reward approach and seeking, while the intermixed population of VP glutamate neurons instead promote avoidance and aversion. Some have thus suggested a functional dichotomy between these VP subpopulations in reward versus threat. We test this hypothesis by asking how inhibiting VP^GABA^ impacts active and passive defensive responses to learned threat cues in the absence of rewards. We taught GAD1:Cre rats with inhibitory VP^GABA^ DREADDs (or control rats) that a metal probe delivers shock, or that a 20sec auditory cue precedes footshocks. These stimuli thereafter elicit active defensive burying, or passive freezing responses, respectively. We found that VP^GABA^ inhibition with CNO markedly increased stimulus-appropriate defensive responses to both types of learned threats, but failed to consistently alter new learning about them—suggesting VP^GABA^ mediates aversive motivation but not memory formation. VP^GABA^ inhibition also altered threat-related c-Fos expression within VP cell populations, and in their efferent target lateral habenula, but not mediodorsal thalamus—pointing to potential underlying circuit mechanisms of defensive responses. Results indicate that VP^GABA^ neurons not only promote reward seeking as previously reported, but that they may also actively inhibit defensive responses to threats that might otherwise compete with reward seeking. This refines our understanding of subcortical valanced motivation circuits, and may suggest new targets for intervening in disorders like addiction and depression.

**Highlights:** -Inhibiting VP^GABA^ increases defenses against threatening stimuli

-Both active (burying) and passive (freezing) responses were enhanced

-Inhibiting VP^GABA^ excites other VP cells, and neurons in downstream LHb

-VP subpopulations may interact to bidirectionally modulate circuits and adaptive behaviors

**Graphical abstract:** 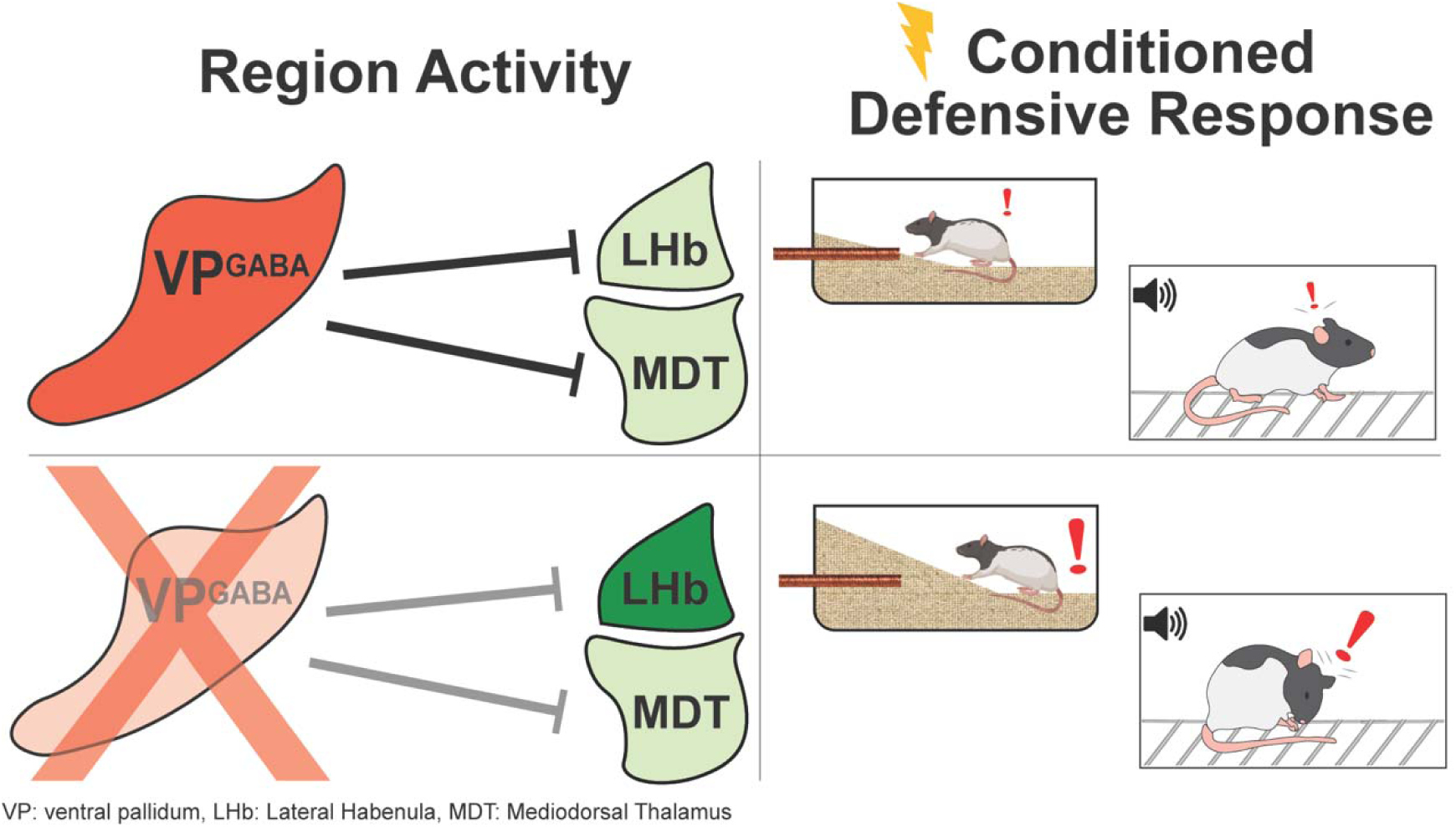

## 1. Introduction

The ventral pallidum (VP) is best known for its role in reward, especially via its strong inputs from nucleus accumbens, and prominent projections to ventral tegmental area (VTA) (Heimer et al., 1991; Kalivas et al., 1993; Mahler et al., 2014; Ottenheimer et al., 2018; Pardo-Garcia et al., 2019; Smith & Berridge, 2007; Zahm, 1989). However, like other nearby hypothalamic and extended amygdala regions (Barker et al., 2017; Jennings et al., 2013a; Lammel et al., 2012; Nieh et al., 2016), VP also contains glutamatergic populations that instead mediate aversion, in part via their direct projections to lateral habenula (LHb) (Haber et al. 1985; Zahm et al. 1996; Faget et al., 2018, 2024; Stephenson-Jones et al., 2020; Tooley et al., 2018; Wang et al., 2020). Yet VP sends both GABA and glutamate projections to LHb as well as reward-related regions (Soares-Cunha & Heinsbroek, 2023)—suggesting unresolved complexity in the specific behavioral roles of these motivation circuits. Since adaptive decision making depends upon proper balance between appetitive and aversive motivations, understanding how their circuit substrates function normally, as well as how they become imbalanced in psychiatric disorders, is essential.

Cell type-specific manipulation and observation studies in mice suggest that VP^GABA^ neurons selectively mediate reward and approach, while VP^glutamate^ neurons instead mediate aversion and avoidance (Heinsbroek et al., 2020; Stephenson-Jones et al., 2020; Tooley et al., 2018; Wulff et al., 2019, Faget et al., 2018, 2024). In rats, we and others find that VP^GABA^ neurons also mediate pursuit of highly salient rewards, especially when triggered by learned reward cues (Farrell et al., 2021, 2022; Scott et al., 2023, 2025; Prasad et. al. 2020). In contrast, we found that inhibiting VP^GABA^ neurons did not alter unconditioned affective responses to unpleasant stimuli like foot shocks in rats, and caused only minor effects on operant avoidance of shock (Farrell et al., 2021, 2022). These data largely support the idea of a highly specific role for VP^GABA^ in appetitive motivation.

This said, we still wondered if VP^GABA^ neurons might play a role in aversive motivation under certain circumstances, especially in response to aversive conditioned stimuli. Might VP^GABA^ neurons be capable of constraining responses to perceived threats in addition to promoting reward seeking, allowing the organism suppress threat responses and to fully commit to capitalizing on opportunities? Since reward-related behavioral effects of VP^GABA^ manipulations are often clearest when motivation is elicited by conditioned cues (Scott et al., 2023; Farrell et. al., 2022), and since few studies had interrogated VP^GABA^ involvement in conditioned *aversive* motivation, we hypothesized that manipulations of VP^GABA^ might alter motivated responses to cues signifying threat, as they do cues paired with reward.

We therefore asked how chemogenetically inhibiting VP^GABA^ neurons affects defensive responses to perceived threats when they are encountered in the absence of reward-related stimuli. We tested both a spatially-localized threat (a probe that can deliver a shock), which normally elicits active defensive burying of the probe, and a temporally-predictive threat (a 20sec auditory cue predicting foot shock), which normally elicits passive defensive freezing. Inhibiting VP^GABA^ neurons augmented context-appropriate defensive responses to both threat cues (be it active defense or passive freezing), without impacting other behaviors, or initial learning about threats. In addition, inhibiting VP^GABA^ neurons increased threat-induced Fos expression in neighboring, likely non-GABAergic (and possibly glutamatergic) VP cells, as well as in the downstream LHb, but not mediodorsal thalamus (MDT)—consistent with selective disinhibition of aversion-related neural circuits.

This report is thus the first to show a clear behavioral role for VP^GABA^ neurons in conditioned aversive motivation, and it supports the idea that VP^GABA^ circuits may not only promote reward seeking directly, but also concurrently suppress threat responses that might otherwise constrain or compete with reward pursuit. We thus provide novel insights into the context-dependent neural mechanisms underlying adaptive responses to salient, affectively-valanced stimuli, and how they may go awry in stress-related disorders.

## 2. Materials and Methods

### 2.1 Subjects

Long Evans male and female GAD1:Cre rats (n=68; n=24 female) and their Cre-negative wildtype (WT) littermates (n=17; n=9 female) were bred in-house, and housed in pairs or trios in ventilated tub cages with corncob bedding and *ad libitum* chow and water. Experiments were conducted during the dark phase of a reversed 12:12hr light cycle. Rats were 75+ days old when experiments commenced. Procedures were approved by the UCI Institutional Animal Care and Use Committee, and followed NIH Guide for the Care and Use of Laboratory Animals guidelines.

### 2.2 Stereotaxic Surgery

GAD1:Cre and WT rats were anesthetized with ketamine (56.5mg/kg), xylazine (8.7mg/kg), and meloxicam (1mg/kg), and an AAV2 vector containing a Cre-dependent, mCherry-tagged inhibitory DREADD (hSyn-DIO-hM4Di-mCherry; titer: 5×10¹² vg/mL; Addgene Cat. #:44362-AAV2) was injected bilaterally with a glass pipette into VP (∼300nl/hemisphere), at bregma-relative coordinates (mm): AP: 0.0, ML: ±2.0, DV: −8.0, as previously described (Martinez et al., 2023). Rats were given at least 21 days to recover from surgery and allow for viral expression to occur before commencing behavioral testing.

### 2.3 CNO Preparation and Injection

Clozapine-N-oxide (CNO) was obtained from the NIDA Drug Supply Program, stored at 4°C in powder aliquots with desiccant, and protected from light. CNO (5mg/kg/ml) was dissolved daily for IP injection in 5% dimethyl sulfoxide, then diluted in 0.9% saline to dose. Vehicle (VEH) or CNO was injected IP 30min prior to commencement of all behavioral tests (Farrell et al., 2019, 2021, 2022; Lawson et al., 2021, 2023; Mahler et al., 2014; Martinez et al., 2024).

### 2.4 Behavioral Methods

#### 2.4.1 Experimental Timeline

At least 4 weeks after surgery, rats were handled for 5min on 5 consecutive days prior to behavioral training. Male and female GAD1:Cre (n=24) and WT rats (n=11) first underwent testing for shock probe fear expression (2 expression tests, counterbalanced VEH and CNO), then they were tested for expression of freezing elicited by 0.75mA foot shock-paired cues in either VP^GABA^-Inhibited (GAD1:Cre+CNO; n=16) or Control-treated (GAD1:Cre+VEH: n=8; or WT+CNO: n=11) rats in a between-subjects design. Other male and female rats first underwent testing for expression of auditory cue fear elicited by 0.3mA foot shock-paired cues after VP^GABA^ inhibition (GAD1:Cre+CNO: n=16) or Control treatment (GAD1:Cre+VEH: n=11; WT+CNO: n=6), then they were subsequently used to test effects of Inhibition (GAD1:Cre+CNO; n=14) or Control treatment (GAD1:Cre+VEH: n=8; or WT+CNO: n=6) on acquisition of shock probe fear learning. Another group of male GAD1:Cre rats expressing VP^GABA^ DREADDs that previously underwent auditory cue fear conditioning training and expression testing (0.5mA shock during training; behavioral data from this pilot experiment not shown) were given 2 weeks off, then they were used to examine neural activity (Fos) elicited by a threatening or non-threatening probe after injection of either VEH or CNO (Threatening Probe + VEH: n=5; Threatening Probe + CNO: n=4; Non-Threatening Probe + VEH: n=5; Non-Threatening Probe + CNO: n=3).

#### 2.4.2 Shock Probe Test General Methods

All shock probe sessions were videorecorded for later quantification of behaviors, as described in **Supplemental Methods**. Rats were first habituated for 30min on two days to a specific plexiglass tub cage filled with corncob bedding (5cm height from cage bottom). They were then returned to the chamber on the following day for a <u>shock probe fear acquisition session</u>. The chamber now had a copper-wrapped plastic probe inserted through a 1cm diameter hole in the wall located 8cm above the cage floor. The probe delivered a 1.5mA shock when touched (Coulbourn Precision Animal Shocker). Rats were placed in the opposite end of the cage from the shock probe, and the shock probe acquisition session continued for 20min after the first rat-initiated shock (Fucich & Morilak, 2018, Berridge et al., 1988; Craft et al., 1988; Pinel & Treit, 1978). The following day, rats underwent a <u>shock probe fear expression session</u>. The chamber was configured identically to the shock probe fear acquisition session, except the probe was inactive and no longer delivered shock when touched.

#### 2.4.3 VP^GABA^ Inhibition Effects on Shock Probe Fear Expression

To test effects of VP^GABA^ inhibition on expression of defensive responses directed at a threatening probe which previously delivered shock, rats received VEH or CNO injection, then were placed into the chamber with the inactive probe for 20min. 48h later, these rats underwent a short (5min) shock probe fear “re-acquisition” session, where the probe was re-electrified, and on which all rats were shocked again at least once. The next day, they underwent a second 20min shock probe fear expression test, conducted identically to the first, other than that they received the alternative injection (VEH/CNO) prior to the session. Rats received VEH and CNO treatments in counterbalanced order in these two shock probe fear expression tests, and defensive burying behaviors were statistically equivalent in rats that received VEH on the first versus the second shock probe fear expression test (Mann-Whitney U=162, p=0.6072). Therefore, data from both shock probe fear expression tests were combined for repeated measures statistical analysis. One rat’s data was excluded from this experiment because it never received a shock during the shock probe fear acquisition session, and 3 rats’ data were excluded because they never emitting treading/digging behavior on either of the two shock probe fear expression tests.

#### 2.4.4 VP^GABA^ Inhibition Effects on Shock Probe Fear Acquisition

To test effects of VP^GABA^ inhibition on initial learning about the shock-delivering probe, rats received either VEH or CNO injections prior to a shock probe fear acquisition session, following prior testing of auditory cue fear expression. 48hrs later, these rats were returned to the chamber with the now-inactive probe, without further injections, for a 20min shock probe fear expression test.

#### 2.4.5 VP^GABA^ Inhibition Effects on Auditory Cue Fear Expression

All sessions were videorecorded for later quantification of behaviors, as described in **Supplemental Methods**. Auditory cue fear conditioning training and testing took place in Med Associates rat operant chambers (30.5x24x21cm; St. Albans, VT), housed within sound-attenuating boxes. Chambers had a peppermint scent (McCormick) during cue/shock training, and standard metal bars that could deliver foot shock composed the chamber floor. On these 8.5min training sessions, rats were placed into the novel chamber, and a house light turned on for a 3min baseline period. Then, a 20sec auditory cue (continuous white noise: 80dB) was played, which was followed immediately by either a low (0.3mA) or high (0.75mA) intensity foot shock, delivered for 2sec. Three cue/shock pairings were made during this training session, with 90sec elapsing between each. 48hrs later, rats received either VEH or CNO injection, then they were placed into a different Med Associates chamber with a distinct metal grid floor, and an orange scent (McCormick) for a 47min auditory cue fear expression session. After a 3min baseline period, the 20sec shock-associated white noise cue was played a total of 24 times, with 90sec elapsing between each. No shocks were delivered during this session.

#### 2.4.6 VP^GABA^ Inhibition Effects on Probe Threat-Elicited Neural Activity

Male GAD1:Cre rats with VP^GABA^ DREADDs were habituated and trained as described in Section 2.4.2 on the shock probe fear acquisition procedure, except that about half the rats underwent the 20min training session with a probe that did not deliver shock. The following day, rats were injected with either VEH or CNO, then they underwent a shock probe fear expression session as described in Section 2.4.2 (probes did not deliver shock on this session). Rats were returned to their home cages for 90min after the session, then they were perfused and brains were prepared for Fos staining and quantification. One rat in the No-Threat+VEH group did not express sufficient mCherry in VP^GABA^ cells for quantification, so it was excluded from mCherry-based VP Fos analyses, but this rat’s data was included in analyses of Fos in VP^ChAT^ neurons, LHb, and MDT, since these analyses do not require identification of VP^GABA^ versus putative VP^nonGABA^ populations.

### 2.6 Histology, Immunohistochemistry, and Microscopy Methods

Details of staining protocols and Fos quantification methods are provided in **Supplemental Methods**.

#### 2.6.1 Preparation of Slices for Staining

All rats were perfused transcardially with chilled 0.9% saline and 4% paraformaldehyde, brains were postfixed in 4% paraformaldehyde for ∼16hrs, cryoprotected in 20% sucrose-azide for 3+ days, then sectioned coronally at 40μm into PBS-azide.

#### 2.6.2 Visualizing & Quantifying VP Virus Expression

Cre-dependent virus expression localized predominantly within VP was verified in each GAD1:Cre rat used in these studies, by analyzing the zone of immunohistochemically-amplified mCherry expression around viral injection sites in relation to substance P-defined borders of the VP (Haber & Nauta, 1983; Zahm & Heimer, 1988, 1990). Lack of mCherry expression was also verified in each WT Control rat.

#### 2.6.3 Visualizing & Quantifying Neural Activity in VP Cell Populations, LHb, and MDT

We used dual color immunofluorescence to stain for Fos protein, and to amplify Cre-dependent mCherry expression in VP^GABA^ cells in the immediate vicinity of the center of virus injection sites. We examined the percentage of VP^GABA^ neurons (i.e. mCherry+ cells, as these were previously confirmed to be ∼91% GABAergic (Farrell et al., 2021)) that also expressed Fos, and we also quantified the number of Fos+ VP cells that did not also express mCherry in the same zone. Although we did not verify the identity of these mCherry-negative VP cells directly, they were well within the zone of Cre-dependent expression of mCherry in VP^GABA^ cells using this highly-penetrant and specific transfection system (Farrell et al., 2021), so we hereafter refer to them as putative VP^nonGABA^ cells. We note that we were unable to directly identify whether these unlabeled cells were VP^glutamate^ using the IHC methods available to us, since vGlut2 protein does not persist long in soma before it is trafficked axonally. Therefore, we further explored the potential identity of these Fos+, putatively VP^nonGABA^ cells by staining additional adjacent sections for Fos and choline acetyltransferase (ChAT), which allowed us to determine whether this cholinergic population was affected by Threat and/or VP^GABA^ Inhibition. Finally, we also Fos-stained sections that contained LHb and MDT from the same animals, allowing us to determine how neural activity in both these VP efferent targets was affected by Threat and/or VP^GABA^ Inhibition, and how this activity relates to activity in VP subpopulations in the same individuals.

### 2.7 Data Analysis Approach

#### 2.7.1 Group Comparisons

To determine effects of VP^GABA^ inhibition on shock probe fear expression, male and female GAD1:Cre and WT rats received VEH and CNO in counterbalanced order before each of two probe fear expression tests, and within-subjects comparisons were used. To determine effects of VP^GABA^ inhibition on shock probe fear acquisition and auditory cue fear expression, GAD1:Cre rats with DREADDs that were injected with CNO (Inhibited Group) were compared to Controls that consisted of a mixture of GAD1:Cre rats with DREADDs that were given VEH (n=8 high-shock intensity experiment, 11 low-shock intensity experiment, 8 shock probe fear acquisition experiment), and WT rats without DREADDs that were given CNO (n=11 high-shock, 6 low-shock, 6 shock probe fear acquisition). No behavioral differences were seen between these subgroups of Control rats in 1^st^ cue freezing, or in defensive burying (ts<1.522, ps>0.05), so they were combined for primary comparisons of Control and Inhibited rats.

#### 2.7.2 Statistical Analyses

Statistical analyses were conducted in Graphpad Prism. ANOVAs and t-tests were used to determine main effects and interactions in normally distributed datasets. When data violated normality assumptions of parametric tests (assessed by D’Agostino & Pearson tests), data were either log-normalized (shock probe fear expression experiment), or non-parametric tests (Mann-Whitney U) were used for independent samples. For auditory cue fear expression tests, a repeated measures Trial (24 Cue Deliveries) factor was included in ANOVAs to examine extinction of cue responses within the session. Secondary analyses for each experiment examined main effects of Sex, and its interactions with other variables. Fos data were analyzed with two-way ANOVA incorporating Drug (VEH, CNO) and Probe Type (Threat, No-Threat), and Pearson correlations examined how Fos in LHb or MDT relates to Fos in VP^GABA^, putative VP^nonGABA^, or VP^ChAT^ subpopulations. All tests were 2-tailed, with significance criterion set at p<0.05.

## 3. Results

### 3.1 Analysis of VP DREADD Expression

The previously-validated (Farrell et al., 2021, 2022), Cre-dependent hM4Di DREADD/mCherry vector caused robust expression in VP^GABA^ neurons of GAD1:Cre rats, but not in their Cre-WT littermates (**Fig. 1A,D**). Somatic mCherry expression in included rats was localized predominantly within Substance P-defined VP borders of GAD1:Cre rats (**Fig. 1B**), as well as in these neurons’ axons, which were present as expected in VP efferent targets including LHb and MDT (Groenewegen & Berendse, 1990; Haber et al., 1993; Root et al., 2015; Zahm et al., 1987) (**Fig. 1C**). Some GAD1:Cre rats were excluded from behavioral analyses for significant DREADD expression observed outside VP borders (n=5), or lack of expected VP mCherry expression (n=9).

**Figure 1:**
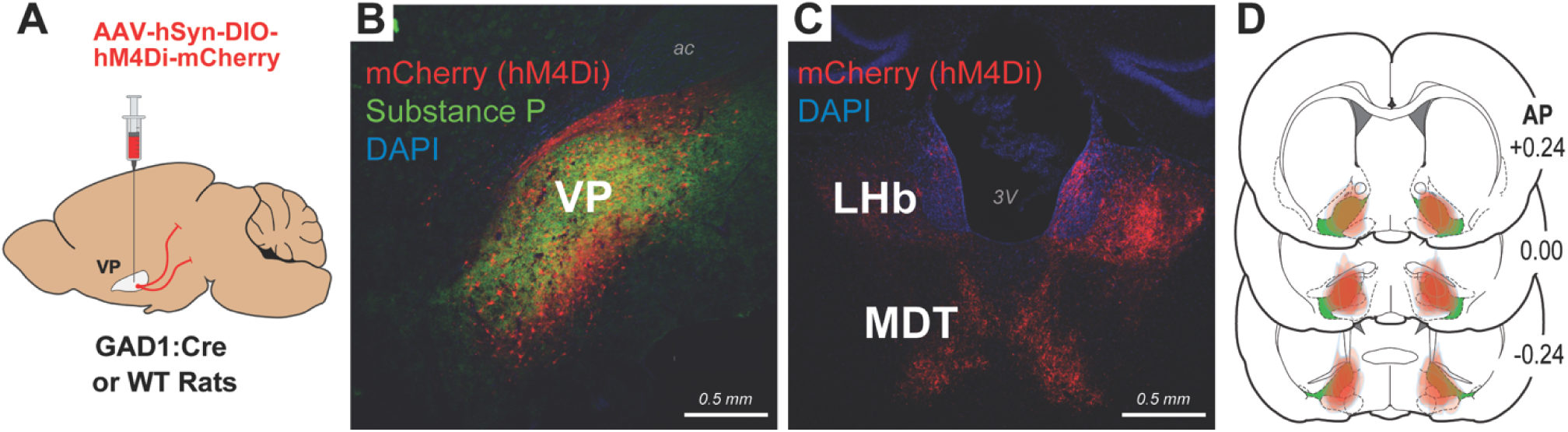
Targeting VP^GABA^ Neurons with Inhibitory DREADDs in GAD1:Cre Rats. **(A)** Diagram of hM4Di-vector injection to the VP. **(B)** Representative image of mCherry (red) expression in the Substance P (green)-defined VP from an included GAD1:Cre rat. DAPI is blue. **(C)** mCherry-expressing VP^GABA^ axons (red) in LHb and MDT are shown, DAPI is blue. **(D)** Location of virus expression in each included GAD1:Cre rat is shown in red, with Substance P-defined VP borders represented in green. Coordinates indicate distance anterior or posterior of bregma, in mm. AP=anterior-posterior coordinate relative to bregma. 3V=3^rd^ ventricle.

### 3.2 VP^GABA^ Inhibition Enhances Active Defensive Responses to a Localized Threat

We first asked whether inhibiting VP^GABA^ neurons impacts defensive burying behaviors directed at a localized, previously-learned threat—a probe that delivered an electric shock during training, but which was inactive during counterbalanced VEH and CNO fear expression tests **(Fig. 2A)**.

**Figure 2:**
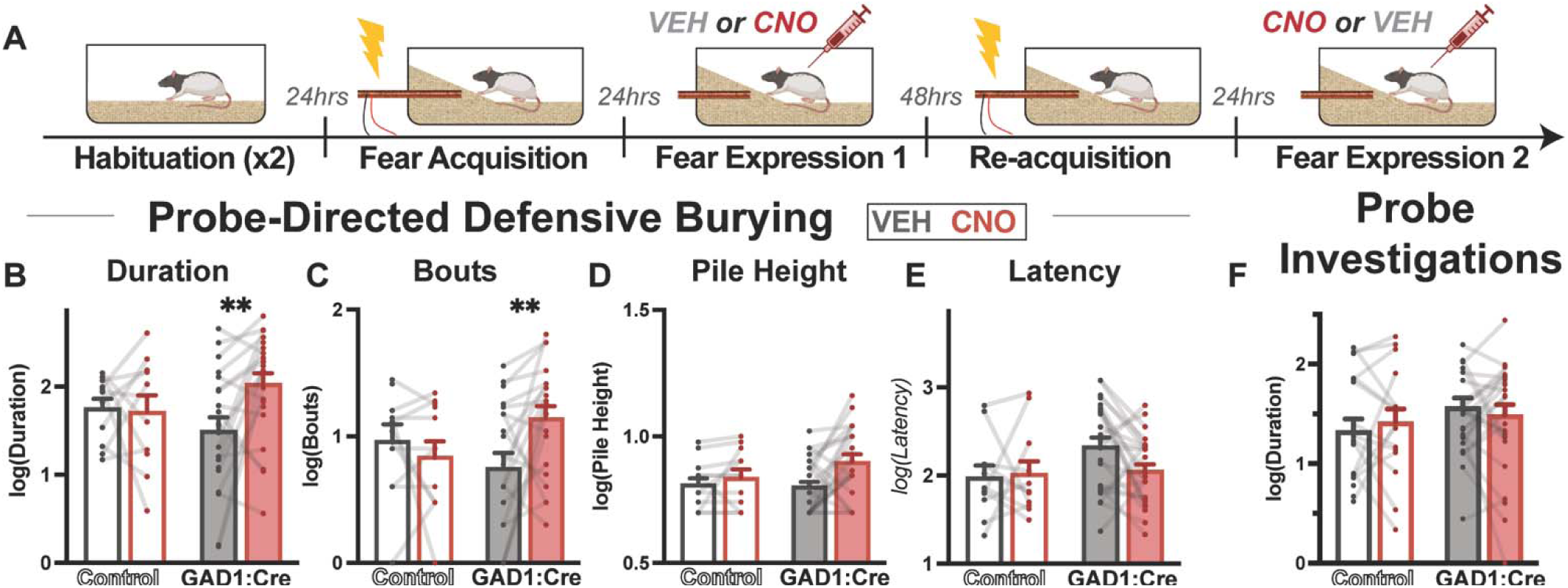
VP^GABA^ Inhibition Enhanced Active Defensive Burying Responses to a Localized Threat. **(A)** Behavioral protocol for probe fear expression tests. **(B-E)** Enhanced defensive burying behaviors in GAD1:Cre rats on CNO day relative to VEH day, with no CNO effect in Control rats. **(F)** In contrast, inhibition caused no change in probe investigations. *p<0.05, Holm-Sidak multiple comparison posthoc. Data were log-normalized to meet ANOVA assumptions, and for display.

#### 3.2.1 Effects of VP^GABA^ Inhibition on Probe Fear Expression

Inhibiting VP^GABA^ neurons markedly increased expression of defensive responses to the previously-learned shock probe. Relative to VEH day, CNO in GAD1:Cre rats (n=11M,13F) increased time animals spent actively burying the shock probe, compared to Controls (n=6F,5M) (DREADD Group x Drug Interaction: F_1,28_=4.196, p=0.050; Holm-Sidak posthoc for VEH versus CNO in GAD1:Cre rats: t_28_=3.226, p=0.007; **Fig. 2B**). Inhibited rats also displayed significantly more bouts of burying behavior (F_1,28_=6.687, p=0.015; Holm-Sidak: t_28_=3.291, p=0.005; **Fig. 2C**) and non-significant trends toward increased height (cm) of bedding piles formed (F_1,32_=2.068, p=0.156, **Fig. 2D**), and decreased latency to begin burying (F_1,32_=2.585, p=0.117, **Fig. 2E**) were observed.

In contrast to these active defensive responses to the probe, inhibiting VP^GABA^ did not alter other behaviors less linked to active defense, such as the number of probe investigations (F_1,32_=0.506, p=0.479, **Fig. 2F**) or escape attempts (F_1,32_=0.27, p=.606). Freezing was uncommon in this active fear expression task and was unaffected by VP^GABA^ inhibition (data not shown).

Effects of CNO in GAD1:Cre rats are attributable to the chemogenetic manipulation of VP^GABA^ neurons, as no significant main effects of Drug were observed in the DREADD Group x Drug analysis for burying bouts (Main effect of Drug: F_1,28_=0.0112, p=0.917) or duration (F_1,28_=3.056, p=0.09). During initial drug-free learning of the shock probe threat, GAD1:Cre and Control rats acquired the task similarly, indicating that observed VP^GABA^ inhibition effects did not result from pre-existing group differences in training history or efficacy. All rats contacted the probe at least once during initial shock training, and the duration and number of bouts of defensive burying did not differ between GAD1:Cre and Control rats (DREADD Group x Drug Interaction: Fs<1.734, ps>0.199; **Fig. S1A-B**). Results thus indicate that inhibiting VP^GABA^ neurons robustly and selectively increased active defensive responses directed at a previously-learned localized threat.

#### 3.2.2 Similar Effects of VP^GABA^ Inhibition on Probe Fear Expression in Both Sexes

Only minor sex differences were found in this probe fear expression test. Overall, males spent slightly more time burying than females (Main effect of Sex, log-normalized data: F_1,31_=5.128, p=0.031), and built taller piles of bedding (F_1,31_=8.620, p=0.006), while females spent more time attempting to escape than males (Main effect of Sex: F_1,37_=32.630, p<0.0001). There were no other sex differences in probe-directed behaviors (burying bouts, latency, pile height, probe investigations) or freezing (Fs<3.636, ps>0.05; **Fig. S2A-D)**.

No sex differences were seen in effects of VP^GABA^ inhibition on defensive responses to the probe, since no significant Sex x Drug interactions were observed for any behavior in Inhibited rats, including burying, pile height, latency to bury, freezing, escape attempts, or probe investigations (all Fs<2.573, ps>0.120; **Fig. S2A-D**). No sex-specific effects of CNO versus VEH were observed in Control animal behaviors either (no Sex x Drug interactions, log-normalized data; Fs<2.572, ps>0.05).

### 3.3 VP^GABA^ Inhibition Does Not Consistently Alter Learning About the Shock Probe

#### 3.3.1 Effects of VP^GABA^ Inhibition on Active Fear Acquisition

Next we asked whether inhibiting VP^GABA^ impacts learning about the shock-delivering probe, by training a cohort of Inhibited (n=15;6F,9M) and Control rats (n=15;6F,9M) to acquire a fear memory during VP^GABA^ manipulation (**Fig. 3A**). VP^GABA^ inhibited rats did not differ from Controls in defensive burying during this acquisition session (Mann-Whitney test; duration: U=85, p=0.563, **Fig. 3B**; bouts: U=65.5, p=0.140, **Fig. 3C**; pile height: U=89, p=0.689, **Fig. 3D**; latency: U=65, p=0.137, **Fig. 3E**). Probe investigations (U=95, p=0.834, **Fig. 3F**) and freezing (U=88, p=0.667) were similarly unaffected during acquisition. Rats were then tested 48hrs later with the non-electrified probe, drug-free, thus querying retention of the threat learning that occurred in the presence or absence of VP^GABA^. Again, no significant differences between Inhibited and Control rats were observed overall in defensive responses (duration: U=87.5, p=0.643, **Fig. 3G**; bouts: U=84.5, p=0.765, **Fig. 3H**; pile height: U=88.5, 0.673, **Fig. 3I**; latency: U=95.5, p=0.918, **Fig. 3J**) or probe investigations (U=62, p=0.104, **Fig. 3K**), and freezing was not observed in any rats. These data suggest that VP^GABA^ inhibition does not consistently alter unconditioned responses to an electrified shock probe itself, nor learning or consolidation of memories about this threat.

**Figure 3:**
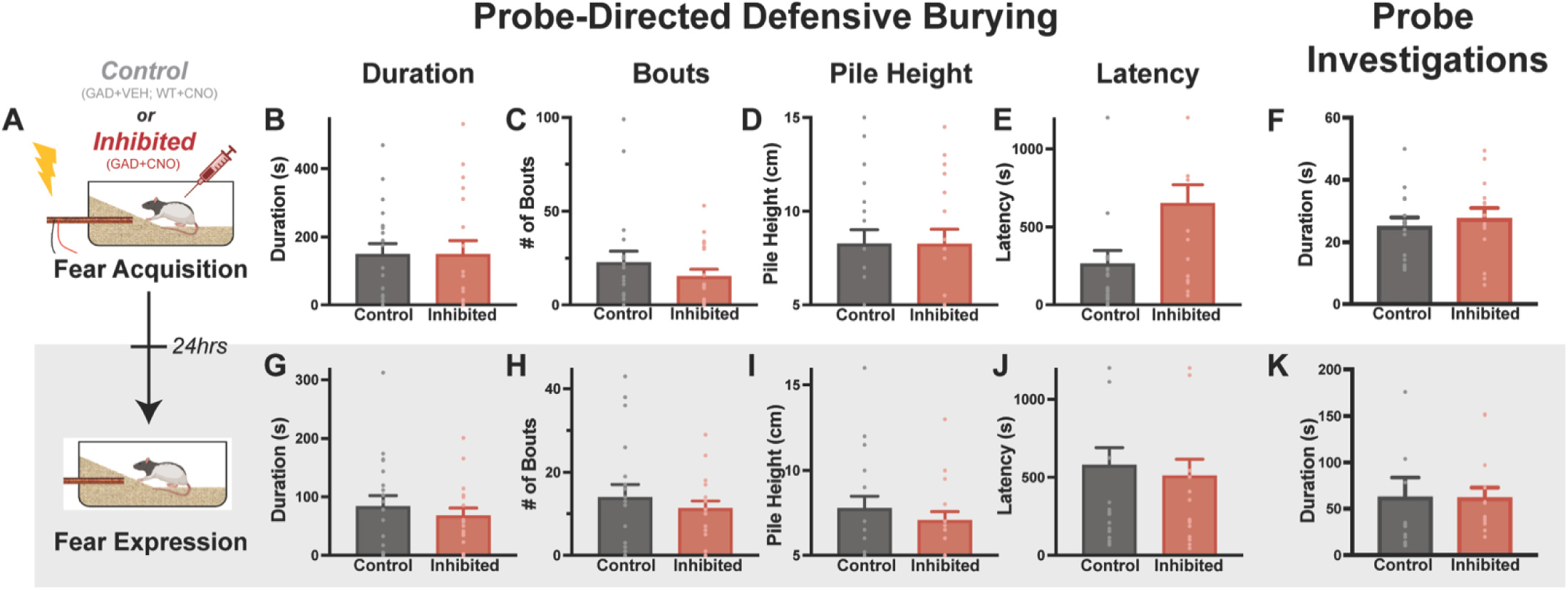
VP^GABA^ Inhibition During Training Does Not Consistently Affect Acquisition of Defensive Burying. (A) Diagram shows behavioral protocol, where CNO or VEH injections were given during shock probe training in Control or Inhibited groups, then a drug-free memory test was conducted with a dormant probe. Training day behaviors are shown for defensive burying duration **(B)**, bouts **(C)**, pile height **(D)**, and latency **(E)**, and also probe investigation duration **(F)**. The same behaviors on the memory test are shown in **(G-K).** No statistically significant results were found.

#### 3.3.2 Sex Differences in Active Fear Acquisition Tests

Female rats displayed slightly less burying behavior than males during shock probe acquisition (main effect of Sex; burying duration: F_1,22_=3.378, p=0.080; bouts: F_1,22_=5.864, p=0.024; pile height: F_1,24_=7.225, p=0.013; **Fig. S3A-C**), and also an increased latency to first bury after being shocked (F_1,24_=6.907, p=0.015; **Fig. S3D**). Accordingly, during the subsequent retention test females were also slower to initiate defensive responses (F_1,24_=8.519, p=0.008, **Fig. S3H**), and they created shorter piles than males (F_1,24_=5.226, p=0.032, **Fig. S3G**).

VP^GABA^ inhibition did not sex-dependently influence defensive behaviors emitted during the acquisition session, as no Sex × DREADD interactions were observed (all Fs<0.600, ps>0.446). However, during the drug-free memory retention test conducted 24hrs later, there was a significant Sex × Inhibition Group interaction, such that prior VP^GABA^ inhibition appeared to sex-dependently affect most measures of defensive behavior (duration: F_1,21_=11.76, p=0.003; bouts: F_1,21_=9.446, p=0.006; pile height: F_1,24_=7.832, p=0.010; latency: F_1,24_=2.789, p=0.108; **Fig. S3E-G**). Posthoc analysis of behavior on this memory retention test in each Sex showed that males which were VP^GABA^-Inhibited during training spent less time burying on the retention test than Control males (Holm-Sidak Posthoc: t_21_=2.471, p=0.044), while females inhibited during training spent more time burying than Control females (t_21_=2.450, p=0.044). Males inhibited during training also had fewer bouts of burying during the retention test (t_21_=2.559, p=0.036), but no effect was seen in females on this measure. No significant differences in pile height during the retention test were seen in females (t_21_= 1.783, p=0.167), while males showed a trend towards a decrease in pile height (t_21_=2.250, p=0.067). Notably, unlike the previous experiment where effects VP^GABA^ inhibition on expression of probe fear were specific to active defensive behaviors, a trend toward a similar sex difference in probe investigations during the memory retention test was also seen (Sex x Inhibition interaction: F_1,24_=3.369, p=0.079). The significance of this potential sex difference in responses to a probe initially encountered during VP^GABA^ inhibition is unclear, so future studies with larger cohorts and complementary behavioral tasks will be important to clarify whether sex mediates VP^GABA^ roles in memory formation or consolidation.

### 3.4 VP^GABA^ Inhibition Enhances Passive Defensive Freezing to a Shock-Predictive Auditory Cue

Next we asked whether VP^GABA^ neurons are also involved in responses to cues that are temporally-but not spatially-predictive of a shock, which elicit passive (freezing) rather than active (probe burying) defensive responses.

Rats first learned to associate a 20sec auditory cue that terminated coincident with a foot shock 3 times during a fear acquisition session (low-shock: 0.3mA, or high-shock: 0.75mA). 48hrs later they underwent a fear expression test in a novel context with VP^GABA^ Inhibition or Control treatment. The shock-paired cue (without shock) was played 24 times, allowing examination of VP^GABA^ inhibition effects on cued fear expression (response to 1^st^ delivered cue) and extinction (subsequent cues; **Fig. 4A)**.

**Figure 4:**
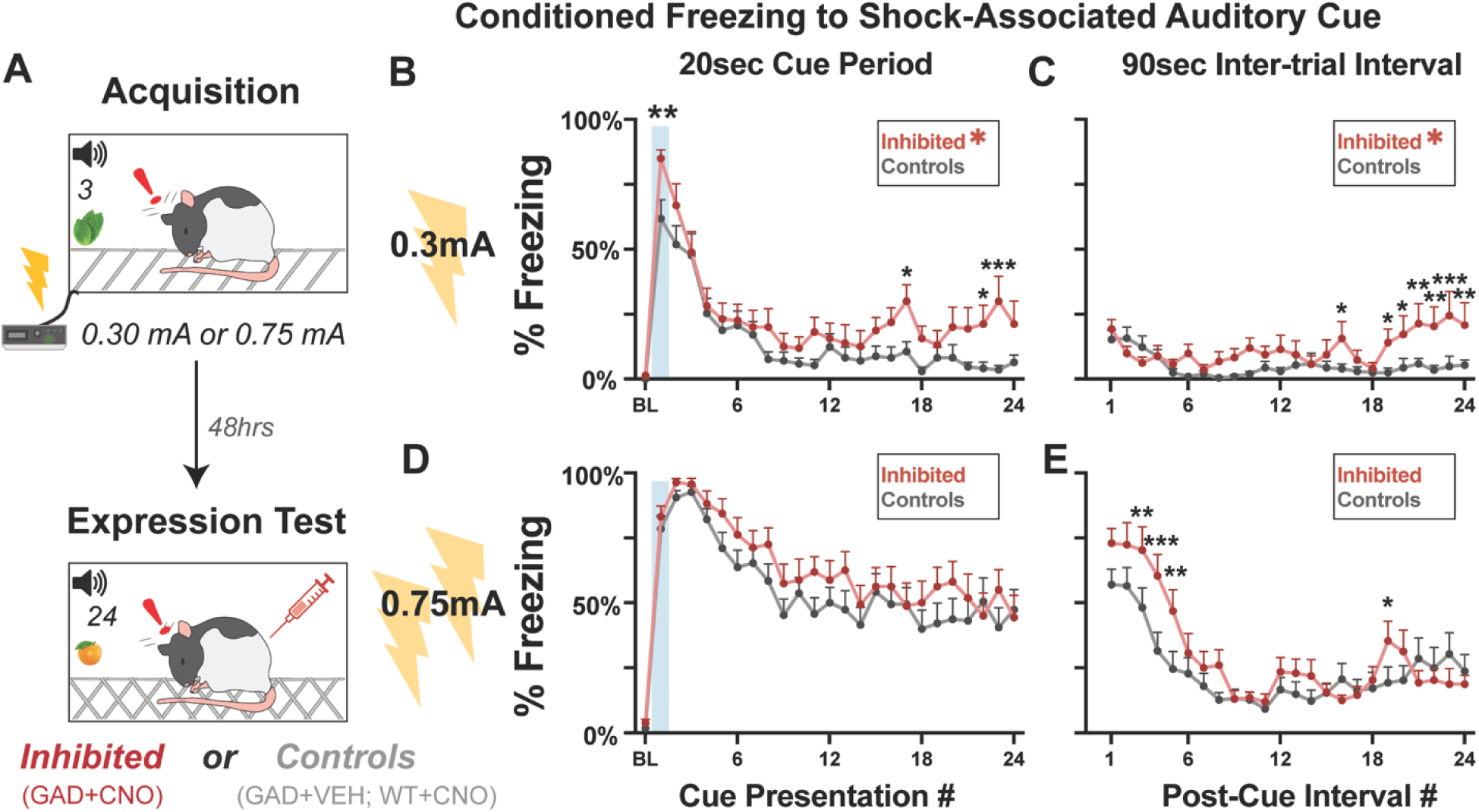
VP^GABA^ Inhibition Enhances Freezing Responses to a Previously-Learned Auditory Fear Cue. **(A)** Diagram of the passive auditory fear expression protocol. During cue/shock Acquisition training, 3 cue/shock (0.3 or 0.75 mA) pairings were made, and 48hrs later, a fear expression test occurred in VP^GABA^-Inhibited and Control rats, where the cue was played 24 times without additional shocks. **(B)** The low-shock intensity (0.3mA) cue yielded enhanced freezing in Inhibited rats relative to Controls during its first presentation (indicated by shaded area), and also across all 24 cue presentations. **(C)** Inhibited rats also froze more than Controls during the 90sec periods following each cue presentation. **(D)** The high-shock intensity (0.75mA) cue elicited similar freezing in Inhibited and Control rats during cue presentations, though **(E)** Inhibited rats froze more during post-cue periods. BL: 3min baseline prior to the first cue. Red asterix = main effect of VP^GABA^ Inhibition (Inhibited versus Control; p<0.05 in a 2-way ANOVA). Black asterix = cue presentations during or after which Inhibited rats froze more than Controls (p<0.05, Holm-Sidak posthoc).

#### 3.4.1 Effects of VP^GABA^ Inhibition on Passive Defensive Responses to Auditory Threat Cues

In the low-shock intensity cohort, VP^GABA^ inhibition (n=8M,9F) did not alter behaviors relative to Control treatment (n=9M,5F) during a baseline period in the novel context (Inhibited versus Control, Mann-Whitney test; U=110.5, p=0.103, **Fig. 4B**), suggesting no change in generalization of fear to the novel chamber as a result of VP^GABA^ inhibition. When the low-intensity shock-paired cue was played for the first time on this expression test, when rats maximally expected to be imminently shocked, Inhibited rats froze more than Controls (U=56.5, p=0.005. **Fig. 4B**, shaded area). Cues were played 23 more times without shocks in the session, and Inhibited rats froze more than Controls during cues throughout this extinction training (Main effect of Inhibition: F_1,31_=6.472, p=0.016; including during the 1^st^, 17^th^, 22^nd^, and 23^rd^ cue presentations; Holm-Šídák postdoc, 1^st^: t_775_=2.976, p=0.003; 17^th^: t_775_=2.486, p=0.013; 22^nd^: t_775_=2.194, p=0.028; 23^rd^: t_775_=3.390, p=0.001). Inhibited and Control groups otherwise showed similar extinction rates (Main effect of Cue#: F_24,744_=23.240, p<0.0001; No Inhibition x Cue# interaction: F_24,744_=1.048, p=0.400).

In addition to measuring behavior during 20sec cues, we also examined the 90sec periods immediately after each cue, and found that VP^GABA^ inhibited rats froze more than Controls across all cues (Main effect of Inhibition: F_1,31_=5.128, p=0.031; **Fig. 4C**), especially late in the session (Inhibition x Cue# interaction: F_24,744_=1.906, p=0.006; Holm-Sidak posthoc significance for cues 16, and 19-24 (ts>2.106; ps<0.036).

For the cohort receiving high-intensity shocks during training, VP^GABA^ inhibition (n=9M,7F) during the expression test did not notably alter behavior relative to Controls (n=9M,10F) during baseline (U=122, p=0.293; **Fig. 4D**), during the first cue presentation (U=196.5, p=0.930; shaded area), or across the 24 cue presentations (no main effect of Inhibition: F_1, 33_= 1.701, p=0.201, nor Inhibition x Cue# interaction: F_24,792_=0.629, p=0.916, **Fig. 4D**), potentially due to a ceiling effect caused by high rates of freezing in both groups. However, during post-cue periods, increased freezing in Inhibited rats was seen in early, but not late extinction trials (Inhibition x Cue# interaction: F_23,759_=2.316, p=0.001, **Fig. 4E**), potentially suggesting that VP^GABA^ inhibition caused “spillover” of fear into these otherwise relatively “safe” post-cue periods. Holm-Sidak posthocs confirmed elevated post-cue freezing during cues 3-5 and 19 (ts>1.981; ps<0.048).

Effects of VP^GABA^ inhibition on expression of auditory cue-elicited freezing did not appear to result from group differences in Pavlovian training, prior to VP^GABA^ inhibition. Inhibited and Control rats behaved similarly during these training sessions, with equivalent freezing during baseline (Us>119, ps>0.05), during each of the three 20sec cue presentations (Us>82.5, ps>0.05), and during 90sec periods following each shock (Us>97.5, ps>0.05; **Fig. S1C,D**).

VP^GABA^ inhibition enhancement of passive defensive responses to previously-learned auditory threat cues was specific to freezing, as other measured behaviors in the low-shock cohort were either unaffected by VP^GABA^ inhibition (rearing, head movements, escape attempts), or they were decreased at the expense of increased freezing. Sniffing was decreased during the 1^st^ cue (U=64, p=0.009), and grooming was decreased during the 24 cues (main effect of Inhibition: F_1,31_=10.06, p=0.004) and the 24 post-cue periods (F_1,31_=7.648, p=0.010). In the high-shock cohort, decreases in nonfreezing behaviors were only seen in the post-cue interval and dependent on both a VP^GABA^ inhibition and time of the post-cue interval during the extinction session (Inhibition x Cue# interaction, rearing: F_23,943_=1.780, p=0.013; head movement: F_23,943_=1.697, p=0.022). We note that defensive burying was never seen in this experiment, since a grid floor was placed above bedding in test chambers.

#### 3.4.2 Sex-Specific Effects of Passive Fear Expression and VP^GABA^ Inhibition

Females froze more than males in general during auditory fear cue experiments, with increased freezing during cues (**Fig S4,** main effect of Sex in low-shock cohort: F_1,28_=7.945, p=0.009; high-shock cohort: F_1,31_=40.820, p<0.0001), and in post-cue periods (low-shock cohort: F_1,28_=9.943, p=0.004; high-shock cohort: F_1,31_=0.597, p=0.446).

VP^GABA^ inhibition similarly enhanced freezing to the 1^st^ cue presentation during recall in the low-shock cohort (Main Effect of Inhibition: F_1,29_=9.003, p=0.006) in both sexes (Sex x Inhibition interaction: F_1,29_=0.224, p=0.639 **Fig. S4A**). However, there was some evidence at VP^GABA^ inhibition had greater ability to augment conditioned freezing to the low intensity shock-paired cue in females, relative to males over the entire extinction session (**Fig. S4**; Sex x Inhibition interaction; low-shock cohort: F_1,28_=4.479, p=0.043; high-shock cohort: F_1,31_=0.754, p=0.392). Similar effects were observed in post-cue periods (low-shock cohort: F_1,29_=5.883, p=0.022; high-shock cohort: F_1,31_=0.962, p=0.334). This intriguing finding is a rare example of a potentially sex-dependent effect of VP^GABA^ inhibition which should be replicated.

### 3.5 VP^GABA^ Inhibition Effects on Neural Activity in LHb, and in Neighboring VP Neurons

#### 3.5.1 Effects of Probe Threat and VP^GABA^ Inhibition on Fos in VP Efferent Targets

We next sought to investigate the neural circuits that may mediate augmented threat responses when VP^GABA^ neurons are inhibited. To do so, we examined Fos expression that was elicited by a threatening probe relative to a probe that when previously encountered did not deliver shock (Threat versus No Threat), and/or by VP^GABA^ inhibition (GAD1:Cre+CNO versus GAD1:Cre+VEH). We quantified Fos in major targets of VP^GABA^ efferent projections (LHb and MDT), and in subpopulations of VP neurons (**Fig. 5A; Fig. S5**).

**Figure 5:**
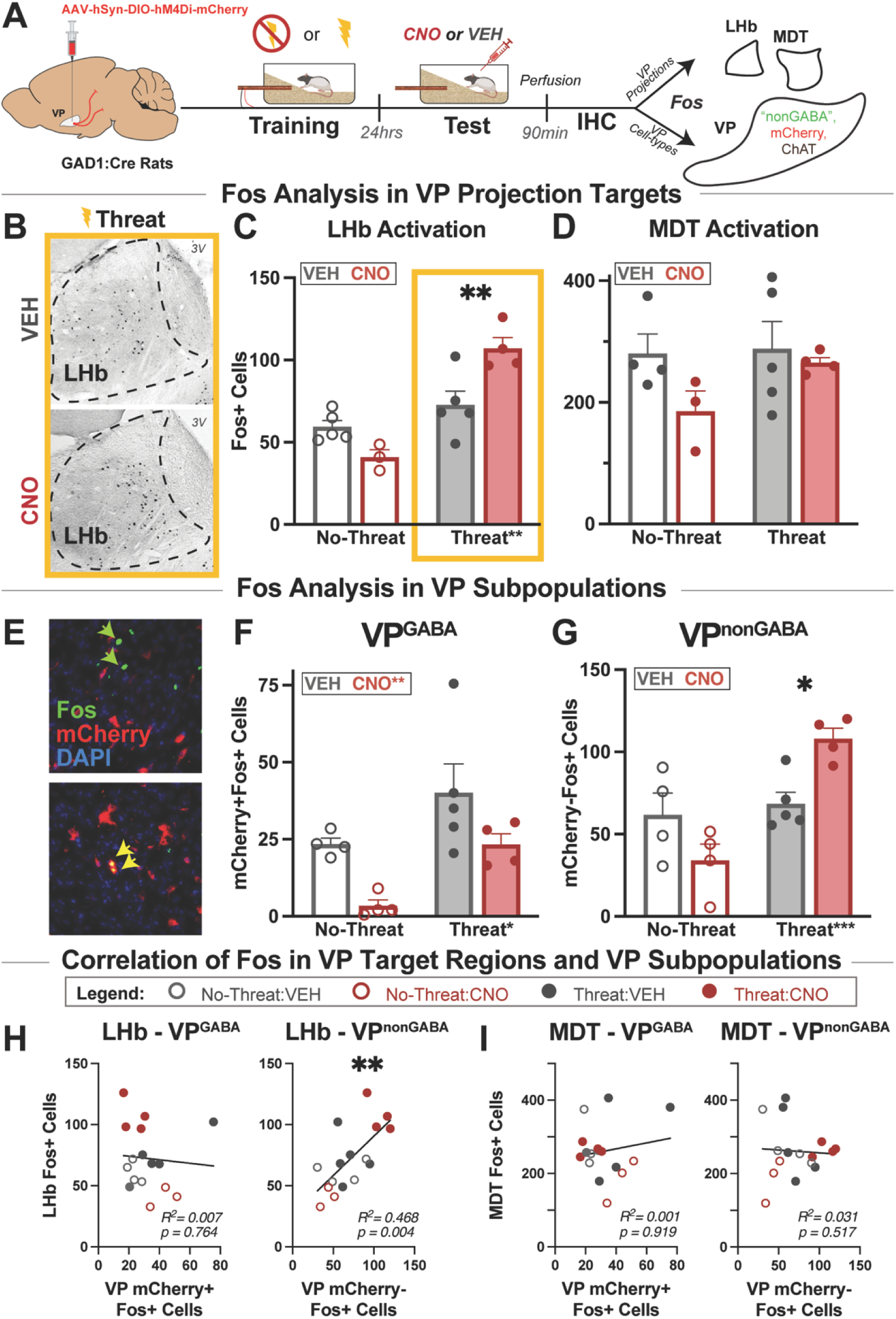
VP^GABA^ Inhibition Increases Threat-induced Fos in the LHb, and in Putative VP^nonGABA^ Cells: **(A)** Schematic of Fos experiment protocol. **(B)** Representative images of LHb Fos staining in Threat+VEH and Threat+CNO conditions, indicated with yellow box. **(C)** Both Threat and VP^GABA^ inhibition increase Fos in LHb, and Inhibition also augmented Threat-induced LHb Fos without affecting LHb Fos in the absence of Threat. **(D)** VP^GABA^ Inhibition and Threat do not significantly impact Fos in MDT. **(E)** Representative images of VP cell populations quantified in the immediate vicinity of virus injection sites in central VP, which were stained for Fos (green, identifying active neurons), mCherry (red, identifying VP^GABA^ neurons), and DAPI (blue, cell nuclei). Yellow arrows indicate active VP^GABA^ cells; green arrows indicate active, putatively VP^nonGABA^cells. **(F)** CNO inhibited mCherry-labeled VP^GABA^ cells, and Threat activated them, but no Threat specific changes were caused by CNO. **(G)** Putative VP^nonGA^ cells were activated by Threat and inhibiting VP^GABA^ cells wit CNO selectively augmented this Threat response. **(H)** Correlation between Fos in LHb and VP^GABA^ (*left*) or putative VP^nonGABA^ (*right*) populations from the same ra are shown. **(I)** Correlation between Fos in MDT and VP^GABA^ (*left*) or putative VP^nonGABA^ (*right*) populations from the same rats are shown 3V=3^rd^ ventricle. Two-Way ANOVA and Holm-Sidak postdoc, or Pearson correlation significance: *p<0.05, **p<0.0 ***p<0.001; two-tailed. Significant main effect of Threat is indicated on X-axis label, and main effect of Drug is shown in legend inset.

We first examined Fos in the LHb, an aversion-related brain region to which VP neurons (including both VP^GABA^ and VP^glutamate^ subpopulations) strongly project (Faget et al., 2018; Groenewegen et al., 1993; Haber et al., 1993; Liu et al., 2020; Root et al., 2014; Tooley et al., 2018), and compared it to another VP target linked more to memory and cognition rather than to aversion, the MDT (Corbit et al., 2003; Floresco et al., 1999b; Leung et al., 2024; Mitchell & Chakraborty, 2013; Ostlund & Balleine, 2008; Parnaudeau et al., 2018; Root et al., 2015; Wolff & Halassa, 2024). Threat (i.e. exposure to a probe that previously delivered shock, or which did not) strongly impacted LHb Fos (2-way ANOVA; main effect of Threat: F_1,13_=33.80, p<0.0001), with VP^GABA^ inhibition significantly enhancing LHb Fos for the shock probe only (Drug x Threat Interaction: F_1,13_=15.01, p=0.002; Holm-Šídák postdoc for VP^GABA^ inhibition effect on response to Threat: t_13_=3.729, p=0.005; or No Threat: t_13_=1.835, p=0.090; **Fig. 5B,C**). These effects appeared to be specific to LHb, as in MDT there were no main effects of Threat (F_1,12_=1.475, p=0.2479), VP^GABA^ inhibition (F_1,12_=2.665, p=0.1285), or interactions of these variables (F_1,12_=0.9897, p=0.3394; **Fig. 5D**).

#### 3.5.2 Effects of Probe Threat and VP^GABA^ Inhibition on Fos in VP Subpopulations

We used our ability to Cre-dependently label VP^GABA^ neurons with mCherry to identify GAD1+ cells within VP (**Fig. 5E**), and to distinguish them from unlabeled cells in the immediate vicinity of virus injections, which are presumably largely non-GABAergic. Non-GABA neurons in the central VP region quantified here are 2-3x more likely to be glutamatergic than cholinergic (Faget et al., 2018; Tooley et al., 2018). Although we did not confirm their identity here, we refer to these unlabeled cells as putative VP^nonGABA^ neurons.

Relative to VEH, CNO significantly decreased activity in VP^GABA^ (mCherry+) cells regardless of Threat (main effect of Drug: F_1,13_=9.128, p=0.0001; no Drug × Threat interaction: F_1,13_=0.071, p=0.7935; **Fig. 5F**), which is consistent with our prior verifications that chemogenetic inhibition in this transgenic system is capable of inhibiting VP^GABA^ neuron activity (Farrell et. al. 2021). We also found that Threat itself elevated activity in VP^GABA^ cells (main effect of Threat: F_1,13_=9.005, p=0.010).

Fos expression in non mCherry-labeled, putative VP^nonGABA^ neurons was also significantly increased by Threat (main effect of Threat: F_1,13_=15.32, p=0.002). However, in the presence of the threatening probe, putative VP^nonGABA^ neurons were further activated by inhibition of their VP^GABA^ neighbors—an effect that was not seen in these neurons in the absence of the shock-associated probe (Drug x Threat Interaction: F_1,12_=10.04, p=0.008; Holm-Šídák postdoc for CNO versus VEH in Threat condition: t_12_=3.278, p=0.013; **Fig. 5G**).

Although we were unable to examine Fos in confirmed VP^glutamate^ neurons, we were able to quantify Fos in VP acetylcholine cells (VP^ChAT^), which comprise the remainder of non-GABA, non-glutamate neurons in VP (Faget et al., 2018; Tooley et al., 2018). VP^GABA^ inhibition (VEH/CNO) did not alter responses of VP^ChAT^ neurons to Threat (t_9_=1.342, p=0.380), nor did VP^ChAT^ Fos differ in either Threat group compared to No Threat rats that received VEH (one way ANOVA: F_2,9_=0.390, p=0.688; Threat+CNO group was not included in this control experiment; **Fig. S5A**). This finding suggests that neither Threat nor VP^GABA^ inhibition appears to alter activity in VP^ChAT^ neurons, suggesting that they are unlikely to account for the clear group differences seen in putative VP^nonGABA^ neurons.

#### 3.5.3 Relationship Between Activity in VP Efferent Targets and VP Subpopulations

We also examined how Fos in VP^GABA^, putative VP^nonGABA^, and VP^ChAT^ populations relates to Fos in VP’s efferent targets LHb and MDT within the same animals, thus pointing toward potential circuit mechanisms that could underlie the observed behavioral effects of VP^GABA^ inhibition in responses to Threat. There was a clear correlation between Fos in putative VP^nonGABA^ cells and LHb (R^2^=0.468, p=0.004, **Fig. 5H, right**), but no such relationship between LHb and VP^GABA^ (R^2^=0.007, p=0.764, **Fig. 5H, left**), or between LHb and VP^ChAT^ (R^2^=0.107, p=0.326, **Fig. S5B**). No such correlation was seen between Fos in MDT and any of these VP subpopulations either (R^2^s<0.031; Ps>0.517; **Fig. 5I, S5C**). These findings are consistent with activity in LHb (but not MDT) being related to activity in non-GABA, non-acetylcholine VP neurons in VP, which could be glutamatergic. However, it is not clear from these data whether VP^GABA^ inhibition yielded an augmented Threat response in LHb via removal of direct LHb inhibitory inputs, via disinhibition of excitatory VP projections to LHb, or via another circuit mechanism.

## 4. Discussion

Here we show for the first time that in addition to promoting reward seeking, VP^GABA^ neurons are also capable of regulating aversively motivated behaviors. Our findings suggest that chemogenetically inhibiting VP^GABA^ neurons “releases the brakes” on context-appropriate defensive responses to learned threats, likely via direct or indirect interactions with LHb. This suggests that VP^GABA^ cells may promote reward pursuit both by imbuing reward-associated stimuli with incentive salience (Berridge & Robinson, 2003; Farrell et al., 2018; Smith et al., 2009), but also by concurrently suppressing conditioned fear and aversion that would otherwise limit reward seeking. We found that VP^GABA^ inhibition consistently enhanced either active or passive defensive responses to shock threat cues, depending on which response was most appropriate for the perceived threat encountered. In contrast, inhibiting VP^GABA^ did not consistently alter acquisition of threat learning, and non-defensive behaviors were unaffected, suggesting a specific role for VP^GABA^ in suppressing fear and aversion elicited by threat cues that were spatially or temporally associated with a noxious shock. These findings lend insight into VP modulation of motivation, and may also reflect broader principles by which intermeshed subcortical circuits of valanced motivation govern affective decision making (Carlezon & Thomas, 2009; Gordon-Fennell & Stuber, 2021; Waraczynski, 2016), and thus inform future strategies for intervening in disorders like addiction and depression.

### 4.1 VP^GABA^ Inhibition Augments Defensive Responses to Learned Threats

VP^GABA^ neurons have been described as primarily helping drive appetitive behavior like highly-motivated reward seeking (Faget et al., 2018, 2024; Prasad et al., 2020; Scott et al., 2023; Soares-Cunha & Heinsbroek, 2023; Stephenson-Jones et al., 2020; Wulff et al., 2019), while intermingled VP^glutamate^ neurons instead seem to mediate aversive and avoidance processes that suppress reward pursuit (Levi et al., 2020; Liu et al., 2020; Tooley et al., 2018; Wang et al., 2020). Other reports have previously found a role for undefined VP cells in fear or aversion (Hernández-Jaramillo et al., 2024; Moaddab et al., 2021; Russell & McDannald, 2024), and one previously showed VP^GABA^ cells respond to noxious stimuli (Faget et al., 2024), but their role in conditioned aversive motivation or defensive behaviors was unknown.

We previously showed that chemogenetically inhibiting VP^GABA^ in rats failed to alter either primary unconditioned responses to shocks, or to markedly alter instrumental avoidance or escape from foot shocks (Farrell et al., 2021), and here we recapitulated the finding that primary behavioral reactions to shock do not require VP^GABA^. Instead, we found an unappreciated role for these neurons in conditioned threat responses, as inactivating them markedly *enhanced* defensive responses to shock-associated cues. This implies that VP^GABA^ neuron activity, in addition to directly promoting appetitive responses, may also engage safety signaling mechanisms that dampen aversive motivation elicited by anticipated dangers (Duvarci, 2024; Luo et al., 2018; Sangha et al., 2020).

In the natural world of rats where opportunities and threats often co-occur, this type of push-pull mechanism would be adaptive in situations in which reward pursuit must be prioritized in the face of danger. Consistent with this idea, when we previously asked how inhibiting VP^GABA^ influences risky decision making, we found a shift to a more conservative strategy—promoting choice of a small but safe reward (one food pellet) over a larger, but potentially dangerous option (two pellets, plus a chance of foot shock)(Farrell et al., 2021). We interpreted this as primarily reflecting suppression of motivation to receive reward when VP^GABA^ was inhibited, but the present data suggest that such decision making may also have been influenced by an augmented perception of threat. We therefore propose that VP^GABA^, and potentially other pallidal, hypothalamic, and extended amygdala GABA circuits with a seemingly similar specific role in promoting reward seeking and consumption (Gordon-Fennell & Stuber, 2021; Jennings et al., 2013; Nieh et al., 2015; Nieh, Vander Weele, et al., 2016; Waraczynski, 2016; Zhang et al., 2021), may in fact perform a more nuanced function than previously appreciated, by balancing conflicting motivational influences to guide high-stakes decision making.

Given the VP’s long-known role in motor activation (Austin & Kalivas, 1990; Churchill et al., 1998; Heimer et al., 1982), it is particularly striking that inhibiting VP^GABA^ led to either active, threat-directed defensive treading, or instead to passive immobility in response to threat—depending on the nature of the shock cue presented. The shock probe used here was a localized stimulus that was spatially paired with shock—analogous to naturalistic threats like a predator chasing a rat down an underground tunnel (Berridge et al., 1988; Craft et al., 1988; Pinel & Treit, 1978). Such proximate, localized threats must either be escaped, or if this is impossible, opposed—for example by pushing and throwing any available material toward the threat, as occurs here when rats attempt to bury the probe in bedding material. VP^GABA^ inhibition selectively augmented this defensive behavior in response to the learned probe threat. In contrast, when a diffuse auditory cue is played just prior to delivery of a foot shock, most rats exhibit freezing behavior when the cue is re-encountered. When VP^GABA^ was inhibited under these circumstances, passive freezing was significantly augmented. Non-defensive behaviors were not altered by VP^GABA^ inhibition in either threat task, nor were defensive behaviors inappropriate to the threat at hand (freezing to the probe, treading to the auditory cue). Therefore, VP^GABA^ inhibition seems to augment aversive motivation in a manner that is upstream of the action selection and generation circuits which determine the best behavioral manifestation of fear under the circumstances. A similar position in threat-processing circuits has been proposed for hypothalamic corticotropin releasing factor cells (Bains et al., 2015; Füzesi et al., 2016), but how exactly these interact with VP, or how either population interacts with downstream mechanisms generating specific defensive actions remains to be determined.

Interestingly, inhibiting VP^GABA^ did not increase freezing in a novel context before cues were presented, suggesting no unusual generalization of fear to this environment, or altered perception of mere novelty as threatening. However, we did observe that VP^GABA^ inhibition caused rats to continue to freeze even after termination of shock cues that were played without delivery of the predicted shock, and to partially oppose extinction of these cues. This suggests that VP^GABA^ inhibition may prolong or intensify cue-elicited fear, thereby interfering with the normal resolution or extinction of the fear response. VP^GABA^ neurons are thus involved in shaping not only the selection of appropriate defensive motor strategies, but also the ability of rats to appropriately terminate defensive states once immediate danger has passed, and to extinguish conditioned fear responses.

Furthermore, we interpret these results to indicate that VP^GABA^ inhibition does not elicit *de novo* fear states in the absence of actual environmental threats. In prior reports, we and others showed that VP^GABA^ inhibition reduces highly motivated, but not less-motivated reward seeking (Farrell et al., 2021, 2022)—if this decreased reward pursuit was secondary to a fear-like state, one would expect all types of reward seeking to be similarly affected. Likewise, inhibiting VP^GABA^ in the absence of threat cues, here or in our prior work, does not elicit either spontaneous freezing in operant boxes, or spontaneous burying in bedding-filled tub cages (Farrell et al., 2021, 2022). We also failed to see activation of LHb by VP^GABA^ inhibition when a non-threatening probe was encountered, but only augmentation of threat-elicited LHb activity. Therefore, we conclude that VP^GABA^ inhibition effects are context- and cue-dependent, and that their behavioral manifestation depends critically upon the affective valence of the circumstances at hand.

### 4.2 VP^GABA^ Inhibition Does Not Alter Defenses Against, or Learning About Unconditioned Threats

We found that VP^GABA^ inhibition did not alter defensive responses directed at an unconditioned and recently experienced threat, as avoidance and burying of a novel, still-electrified probe was unaffected. This suggests that VP^GABA^ does not mediate affective or sensory aspects of the unconditioned experience of punishment as previously reported (Farrell et al., 2021), and it demonstrates that rats are capable of mounting acute, active defenses in the absence of VP^GABA^. In addition, subsequent conditioned responses to the shock probe which was encountered while VP^GABA^ was inactivated were also largely unaltered. This suggests that disrupting VP^GABA^ does not consistently affect the ability to form or consolidate aversive Pavlovian memories. There is some evidence that VP plays a role in other types of learning (Dusa et al., 2022; Floresco et al., 1999a; Kaplan et al., 2020; Leung & Balleine, 2013; Macpherson et al., 2019; Péczely et al., 2014; Roman-Ortiz et al., 2021; Stephenson-Jones, 2019; Stephenson-Jones et al., 2020), though most of these studies did not disambiguate motivational from learning processes, or VP cell types. One report showed that optogenetically stimulating VP^GABA^ suppressed auditory cue-induced freezing without affecting acquisition of new fear learning (Roman-Ortiz et al., 2021), consistent with the present result. However, another showed that stimulating predominantly GABAergic VP enkephalin neurons disrupts formation of an inhibitory avoidance memory (Macpherson et al., 2019). Clearly, further work is required to fully understand the conditions in which VP^GABA^ is implicated in the complex processes involved in forming, consolidating, storing, and retrieving the many different types of memory (McGaugh, 2003; Sherry & Schacter, 1987; Squire, 2004; White & McDonald, 2002).

### 4.3 VP^GABA^ Inhibition “Releases Brakes” on LHb, and on Nearby “Non-GABA” Neurons?

The LHb, which helps encode negative affect and learning, and plays a role in aversion, avoidance, and anhedonia (Bromberg-Martin & Hikosaka, 2011; Li et al., 2013; Liu et al., 2020; Matsumoto & Hikosaka, 2009; Tooley et al., 2018; Y. Yang et al., 2018), is a major efferent target of VP^GABA^ as well as VP^glutamate^ neurons (Faget et al., 2018; Liu et al., 2020; Stephenson-Jones et al., 2020; Tooley et al., 2018). We therefore determined how neural activity in LHb, the MDT (another target of VP efferents), and in VP neuronal subpopulations is affected by exposure to a threatening shock prod, and how inhibiting VP^GABA^ cells affects this activity.

As expected, encountering a threatening probe increased activity in LHb relative to an inert probe, and we found that this activation was further potentiated by inhibiting VP^GABA^. This is consistent with a disinhibition mechanism, whereby disrupting VP^GABA^ removes this direct inhibitory input to LHb, amplifying its response to threat. Accordingly, globally inhibiting VP^GABA^ neurons induces place avoidance (Faget et al., 2018), and activating other LHb GABA inputs induces place preference (Barker et al., 2017; Stamatakis et al., 2013). However, optogenetic activation of VP^GABA^ terminals in LHb failed to promote appetitive behavior (Faget et al., 2018), and optogenetic inhibition of this pathway strongly suppresses conditioned reward-seeking, and modestly reduces conditioned avoidance of a noxious stimulus (Stephenson-Jones et al., 2020). Here, we found that VP^GABA^ inhibition in the absence of a threat was not sufficient to activate LHb, suggesting that VP^GABA^ inhibition alone does not recruit LHb, but merely disinhibits its activity when it is recruited by relevant environmental threats. Interestingly, no clear effects of either threat or of VP^GABA^ inhibition were seen in MDT, another major target of VP thought to play a role in memory and cognition, rather than aversion and defensive responding.

Since we did not use pathway-specific approaches to examine functions of VP^GABA^ projections to LHb, we cannot establish whether VP^GABA^ inhibition impacts LHb via its direct projections, or via indirect network mechanisms. Indeed, our data raise the intriguing, though untested, possibility that one way in which VP^GABA^ inhibition could indirectly impact LHb is via disinhibition of nearby populations of VP^glutamate^ neurons that also project there. VP^glutamate^ projections to LHb mediate aversion- and avoidance-related processes (Faget et al., 2018; Heinsbroek et al., 2020; Stephenson-Jones et al., 2020; Tooley et al., 2018; Wulff et al., 2019), and VP^GABA^ cells directly inhibit their glutamatergic neighbors (Faget et al., 2018; Levi et al., 2020; Tripathi et al., 2013). It is therefore possible that the activation of putative VP^nonGABA^ cells that we showed occurred after VP^GABA^ inhibition, and which we found were unlikely to be cholinergic, reflects disinhibition of local VP^glutamate^, cells that project to LHb. This is also consistent with the correlation we saw between Fos in putative VP^nonGABA^ and LHb, but not MDT. However, we were unable to identify VP^glutamate^ cells here directly, and we cannot exclude the possibility that some of these cells are unlabeled VP^GABA^ cells, so this hypothesis must be tested in future using appropriate methods like RNA-based cell identification, vGlut2-based cell- and pathway-targeting approaches, or physiological techniques.

### 4.4 Sex Differences in VP^GABA^ Functions

VP (DiBenedictis et al., 2020; Lee et al., 2024; Scott et al., 2025; L. Yang et al., 2024), and the circuits in which it is embedded (Becker & Chartoff, 2019; Lim et al., 2004; Trainor, 2011; Zachry et al., 2021), appear to differ based on sex; as does the prevalence of symptoms defining many psychiatric disorders these circuits contribute to (Bangasser & Cuarenta, 2021; Cosgrove et al., 2007; Green et al., 2018; Kessler, 2003; Trainor, 2011; Young & Pfaff, 2014). We noticed VP-independent sex differences in threat response, such that females overall exhibited more passive freezing to the associated auditory cue presentation, while males exhibited more active burying of the probe cue. We found only subtle sex differences in the effects of VP^GABA^ inhibition on threat responding, in that VP^GABA^ inhibited females froze more than males to auditory fear cues and post-cue intervals throughout the extinction session, while initial enhanced freezing to the 1^st^ cue under inhibition was independent of sex. In this context it is surprising that our lab’s work in VP has rarely observed sex differences in the effects of VP manipulations on drug and natural reward seeking (Farrell et al., 2019, 2021, 2022; Mahler et al., 2014).

Another sex difference we did observe in the effects of VP^GABA^ inhibition stems from our experiment in which VP^GABA^ was inhibited during initial learning about a shock probe, and subsequent testing of learned responses to that probe in the absence of further VP manipulation. VP^GABA^ inhibition did not alter responses in either sex to the shock-delivering probe during their initial encounters with it, and defensive and avoidance behaviors in the minutes following shock were unaffected. However, when rats’ ability to remember this threat learned under VP^GABA^ inhibition was queried, a possible sex difference emerged: enhanced defensive burying in females, and decreased burying in males, consistent with a sex difference in VP^GABA^ participation in memory consolidation. This is an exciting possibility, but given the low sample sizes involved, and lack of converging evidence from other behaviors in these experiments, we suggest caution in interpreting this finding. Future studies with larger sample sizes, complementary behavioral tasks, and designs capable of dissociating memory consolidation from performance (McGaugh, 1973) will be required to resolve the conditions in which sex modulates VP roles in negative emotion, learning, and/or memory (Blanchard et al., 1991).

### 4.5 Caveats and Limitations

These studies have additional limitations that should be noted, and potentially resolved with further research. We limited the aversive stimuli tested here to shock delivered two different ways—it would be valuable to learn whether VP^GABA^ inhibition also augments other responses to learned (e.g. context fear conditioning) or innate threats (e.g predator odors or looming stimuli). Likewise, naturalistic threat responding also depends critically on factors like threat imminence or proximity which could be manipulated in follow-up studies, and other potential responses to threats like escape or defensive aggression were not possible here, so it remains to be seen how VP^GABA^ helps prioritize such strategies. In addition, VP^GABA^ inhibition here enhanced freezing only to a low-intensity shock cue, which likely reflects a ceiling effect for freezing with the high-shock intensity cue, but could also suggest more complex interactions. Regarding neural activity data, we cannot exclude the possibility that CNO had DREADD-independent effects on LHb Fos, since we did not include a DREADD-free control group in this experiment. However, nonspecific CNO effects are unlikely to account at least for enhanced LHb Fos in VP^GABA^ inhibited rats exposed to threat, since CNO did not augment Fos in unthreatened rats. In addition, Fos analyses of rats exposed to threat were performed in males only, and should be followed up with females, especially since as noted above the nature of sex differences in VP need further clarification more generally. Although these experiments targeted DREADDs to a region of VP spanning its rostrocaudal axis, prior reports have shown functional distinctions between these subregions (Farrell et al., 2022; Ho & Berridge, 2014; Johnson et al., 1993; Kupchik & Kalivas, 2013; Mahler et al., 2014; Panagis et al., 1995; Root et al., 2013; Smith & Berridge, 2005; Stout et al., 2016; Zahm & Heimer, 1993) so future work should test whether such dissociations may also relate to VP^GABA^ roles in aversive motivation.

### 4.6 Conclusions

Here we report a novel role for VP^GABA^ neurons in suppressing responses to threat cues, complementing their established role in promoting reward seeking. These cells may therefore facilitate commitment to reward pursuit even in the face of danger, thus prevent vacillation between behavioral strategies once a decision to pursue reward is made. In addition to clarifying roles for VP circuits in motivation, these findings might also be relevant to the organization of wider subcortical circuits involving seemingly analogous intermixed populations of GABA and glutamate cells that mediate reward and aversion, respectively (Gordon-Fennell & Stuber, 2021; Hu, 2016; Root et al., 2015, 2020; Soares-Cunha & Heinsbroek, 2023; Wulff et al., 2019). These data also open new avenues of investigation for understanding how VP^GABA^ interacts with other VP cell populations involved in stress (Chang & Grace, 2014; Eckenwiler et al., 2025; Ji et al., 2022), and with wider brain circuits of motivation, emotion, and learning. Given that imbalance in motivational drives is likely involved in numerous psychiatric disorders including addiction, depression, schizophrenia, and PTSD, understanding the functions of these brain networks will be essential for developing next-generation strategies for treating these devastating conditions.

## 5. Authorship Contribution Statement

EMR and MXM equally contributed to conceptualization and implementation of the project, and with SVM analyzed data and prepared the manuscript. CMR, RKR, MRF, VAV, SAW, HRR, and GJK contributed data to the project.

## 6. Acknowledgements

This work was supported by the following grants: NIDA R01DA055849, NIMH R01MH132680, NIDA U01DA053826, NIDA P50DA044118-05A1, TRDRP T31IR1767; EMR Ford Foundation Predoctoral Fellowship; MXM NSF GRFP DGE-1839285 and NINDS F99NS141399. We thank Yiyan Xie and Erik Castillo for technical support, Tom Hnasko for clarifying conversations and technical assistance, and NIDA/NIMH drug distribution programs for supplying CNO.

