## Supplemental Methods and Figures for "Ventral Pallidum GABA Neuron Inhibition Augments Context-Appropriate Defensive Responses to Learned Threat Cues"

1.1 ***Behavioral Quantification Methods***

*1.1.1 Shock Probe Fear Tasks:* Side-view video recordings of shock probe acquisition and expression tests were scored by blinded observers. Instances of, and total duration of Defensive Burying was of primary interest, defined as probe-directed forepaw movements directing bedding forward toward the probe, or backward toward the rat’s tail (Berridge et al., 1988; Craft et al., 1988; Pinel & Treit, 1978). Other scored behaviors were: Duration of Freezing*:* sustained immobility other than breathing, not including sleeping postures, Duration of Attempted Escape: when rats attempted to reach the top of the cage and escape, and bouts of Probe Investigations*:* nose pointed at, and within 1cm of the probe without touching it. After rats were removed from the chamber at the end of the session, the Height of Bedding near the shock probe was measured with a ruler as another index of probe burying. Latency to first be shocked on shock probe fear acquisition sessions, and Latency to begin treading at the probe on shock probe fear expression sessions was also recorded.

*1.1.2 Auditory Cue-Elicited Fear Task:* Videos from the auditory cue fear acquisition and expression tests were manually scored by blinded observers using a previously described sampling method (Fanselow, 1979, 1980). During the 3min baseline, the predominant behavior exhibited by the rat during every 8^th^ second was identified and quantified, for a total of 23 samples per baseline period. During each 20sec cue presentation, and for the 20sec period after each cue presentation, behavior was evaluated likewise, but every 2sec, for a total of 10 samples per cue period and post-cue period. The following behaviors were recorded: Freezing was identified by lack of movement other than breathing, when not laying in a sleeping posture. Locomotion was defined as the animal in motion using all four limbs. Head Movement was defined as the animal moving only its head in any direction without visible sniffing behavior. Sniffing was defined as rapid, rhythmic respiration using the nostrils. Grooming was defined when the animal used its paws or tongue to scratch or lick its own body. Escape Attempts were scored when an animal jumped off the floor of the chamber. Primary analyses were conducted on the percentage of collected samples during periods of interest (baseline periods, during each 20sec cue, 20sec following each cue) in which a behavior occurred.

1.2 ***Histology, Immunohistochemistry, and Microscopy Details***

*1.2.1 Visualizing & Quantifying VP Virus Expression:* Following perfusion and slice preparation, VP-containing sections were blocked in PBS with 0.3% Triton-X (PBST), and 3% normal donkey serum (NDS, Vector Laboratories, Burlingame, CA) for 2hr. Sections were incubated for 16hr at RT in 3% NDS PBST-azide with rabbit anti-substance P (ImmunoStar, Cat. #20064; 1:5000) and mouse anti-DSRed (blabelinglabeling Cre-dependent mCherry expression; Takara, Cat. #632543; 1:2000) primary antibodies. After washing, slices were next incubated in the dark at RT for 5hrs in AlexaFluor-donkey anti-rabbit 488 (Invitrogen, Cat. #A21206; 1:500), and AlexaFluor-donkey anti-mouse 594 (Invitrogen, Cat. #A21202; 1:500) in PBST. They were then washed in PB, mounted on slides, and coverslipped with Fluoromount (Thermo Fisher Scientific). Sections were imaged at 5x magnification with a Leica DM4000 with StereoInvestigator software (MBF Bioscience, Williston, VT) and viral spread was noted inside or outside of VP borders as described in section 2.6.2 of the main text.

*1.2.2 Visualizing & Quantifying Fos in VP^GABA^ and Putative VP^nonGABA^ cells:* Sections near the center of mCherry expression within VP were collected from each rat. They were blocked and incubated in primary antibodies (rabbit anti-Fos: Abcam, Cat. #ab190289; 1:5000; mouse anti-mCherry: Takara, Cat. #632543, 1:2000) for 16hrs at RT, then washed and incubated in dark at RT for 5hrs in Alexafluor-donkey anti-rabbit 488 and Alexafluor-donkey anti-mouse 494 secondary antibodies, as described above.

We imaged Fos- and mCherry-stained VP sections at 20X magnification collecting z-stack images (6 images/20um range) centered on the area of maximal VP viral expression in each hemisphere, with 2 slices/rat quantified bilaterally. Images were flattened in the z dimension using the Max Projection tool in Stereoinvestigator software (MBF Bioscience, Williston, VT), each sample was analyzed by a blinded observer offline. Cells expressing mCherry in soma without nuclear Fos were considered inactive VP^GABA^ cells. Those expressing somatic mCherry and nuclear Fos were considered active VP^GABA^ cells. Those expressing nuclear Fos without somatic mCherry were considered active putative VP^nonGABA^ cells. The percentage of VP^GABA^ cells that also expressed Fos, and the number of Fos+ cells without mCherry were of primary interest, and all 4 samples per rat were averaged for group-based analyses.

*1.2.3 Visualizing & Quantifying Fos in VP^ChAT^ Cells:* We collected sections adjacent to those analyzed for Fos+/- mCherry, near the center of observed virus expression in each rat. They were blocked in 1% H_2_0_2_ in PBS for 15 min, then 2% NDS PBST for 2hr, and then incubated for 16hr at RT in PBST-azide with 2% NDS and rabbit anti-Fos primary antibody (Abcam, Cat. #ab190289; 1:5000). After washing, slices were incubated in biotinylated donkey anti-rabbit (Jackson Laboratories, Cat. # 711-065-152; 1:500) for 2hrs, washed, then placed into freshly-mixed Avidin–Biotin Complex (ABC) solution (Vector Laboratories) for 90 min. Nuclear Fos was then visualized in blue/black using DAB solution in Tris buffer, prepared with 0.01% H_2_0_2_ and 0.04% Nickel Ammonium Chloride (Vector Laboratories). Sections were washed and rested for 45min at RT, then incubated in monoclonal mouse anti-ChAT (Millipore Sigma, Cat. # MAB-03831, 1:5000) in PBST-Az at RT for 16hrs. They were washed, incubated in biotinylated donkey anti-mouse secondary (Jackson Laboratories, Ca. # 715-065-150; 1:500) for 2h, then ABC for 90min. Somatic ChAT was visualized in brown with DAB in Tris buffer, prepared with 0.01% H_2_0_2_. After washing and mounting on subbed slides, tissue was dehydrated in alcohol baths and xylenes, and coverslipped in Permount (Fisher Scientific, Hampton, NH).

To quantify activity in VP ChAT+ cells, we brightfield-imaged VP at 10X magnification, and all ChAT+ cells found within estimated VP borders (Paxinos & Watson, 2006) were manually counted on two bilateral sections per rat by a blinded observer. The percentage of ChAT+ cells that expressed Fos was quantified in each sample, all 4 samples per rat were averaged, and per-rat averages were statistically analyzed.

*1.2.4 Visualizing & Quantifying Fos in LhB and MDT:* Sections containing LHb and MDT were collected for each rat, and stained for Fos using the ABC-amplified DAB method described above (these sections were not co-stained for ChAT). We imaged LHb and MDT bilaterally in 3 sections/rat (~2.9mm-4.1mm caudal of bregma) at 5X magnification. The number of Fos+ nuclei in each sample was quantified by a blinded observer using the Photoshop Count Tool (Adobe), and per-rat averages were used to analyze group differences. Initially, we conducted this experiment in Non-Threatening Probe + VEH, Threatening Probe + VEH, and Threatening Probe + CNO groups only, and quantified Fos only in LHb. Following initial peer review, we added a fourth group (Non-Threatening Probe + CNO) to the experiment. Although this last group was stained, imaged and analyzed using identical methods and reagents to the initial run, we included additional sections from previously-stained rats in the added staining run, and verified that Fos staining quality, intensity, and patterns of activity across groups was equivalent between the staining runs before combining data for analyses. We also added additional analysis of MDT Fos after initial peer review, which we quantified in the same sections used to analyze LHb Fos, using the same methods.

**Supplemental Figures**

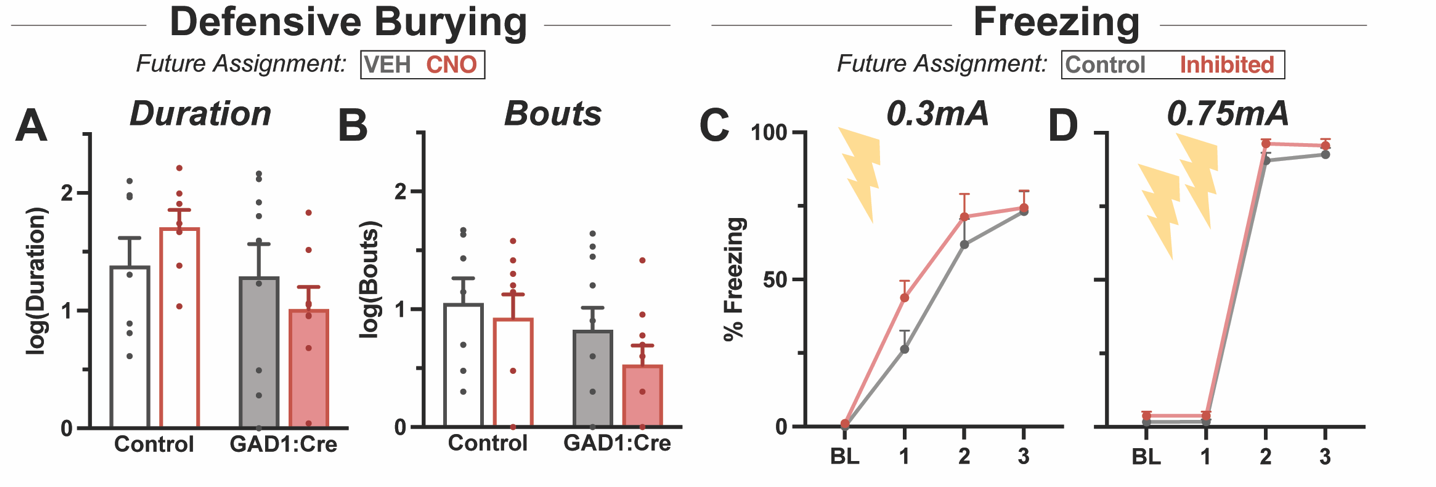

**Supplemental Figure 1: Acquisition of Defensive Behaviors is Similar Prior to Tests of Fear Expression Conducted Under VP^GABA^ Inhibition:** Behaviors during initial acquisition of fear, prior to VP^GABA^ inhibition during subsequent fear expression tests, are shown for shock probe (A-B), and auditory fear conditioning (C-D) experiments. (A) shows duration and (B) bouts of burying in Control and GAD1:Cre rats in training sessions prior to Veh or CNO expression tests. Acquisition of the cue-shock association for low-shock (C) and high-shock (D) cohorts during baseline and the three cues preceding shocks is likewise displayed for rats that would be Control or Inhibited during subsequent fear expression tests.

**Supplemental Figure 2: VP^GABA^ Inhibition Augments Expression of Probe Fear Similarly in Both Sexes.** Burying (A) duration (B) bouts (C) pile height and (D) latency in GAD1:Cre rats is shown in each sex for the probe fear expression experiment. Log-normalized data were used in 2-way ANOVA analysis. Green asterix indicates main effect of Sex, p<0.05.

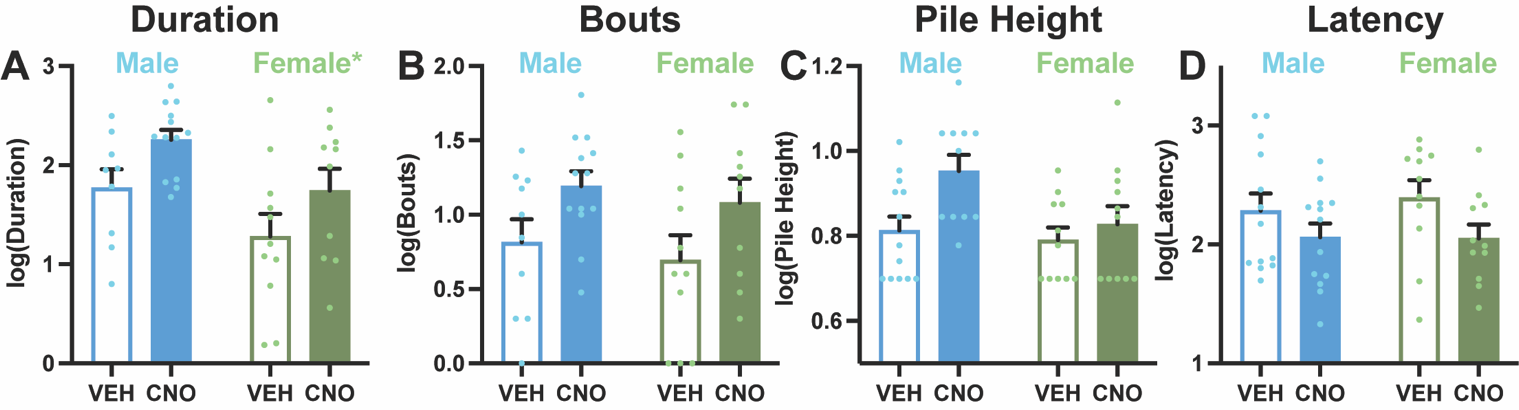

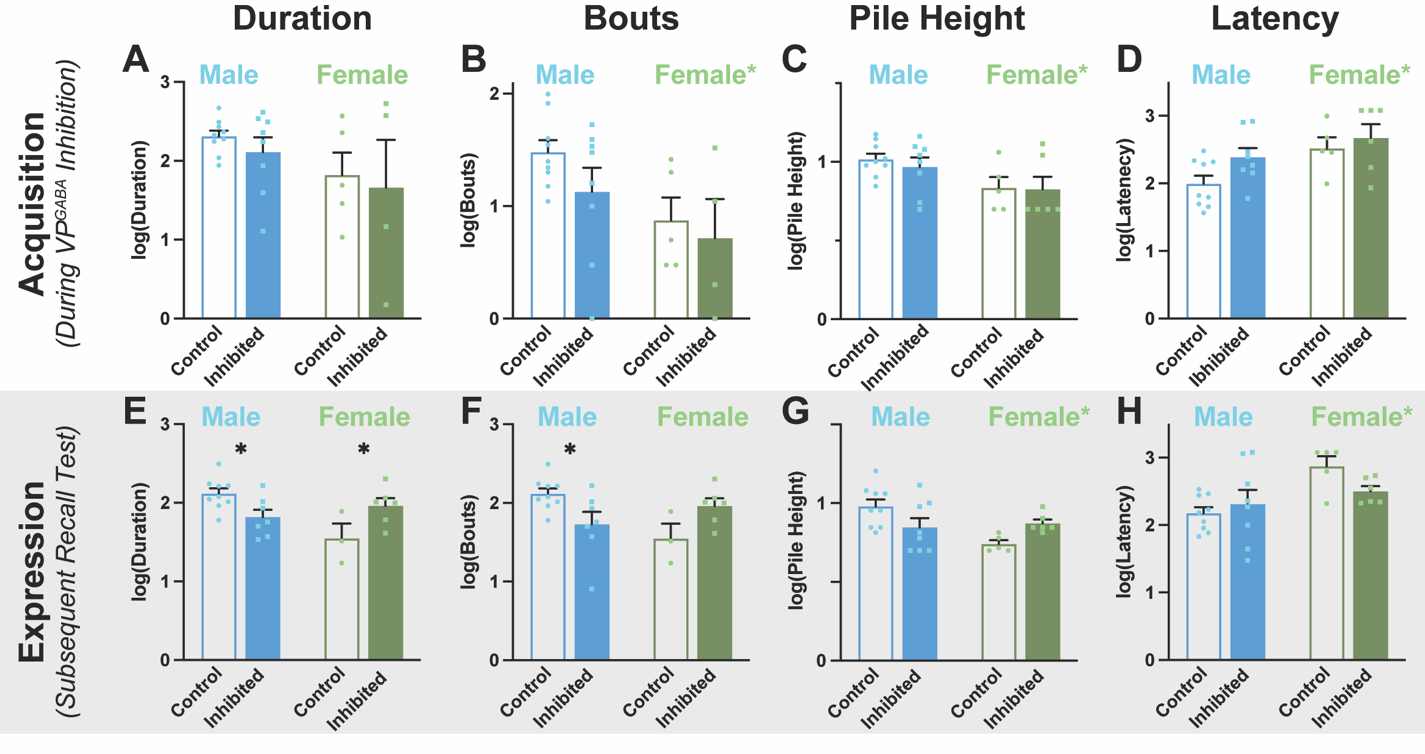

**Supplemental Figure 3: Probe-Directed Behaviors During Initial Shock Training When VP^GABA^ Was Inhibited, and Later During Memory Recall Test, In Each Sex:** Burying (A) duration (B) bouts (C) pile height and (D) latency after shock during probe fear acquisition test when VP^GABA^ was inhibited, shown separately in each sex. 48hrs later these behaviors in the same rats, now with VP^GABA^ intact, are shown in a recall test (E-H). Log-normalized data, 2-way ANOVA of Sex x Inhibition for each behavior. Main effect of Sex indicated with green asterix, black asterix indicates Holm-Sidak posthoc for Inhibition in that sex. p<0.05, two-tailed.

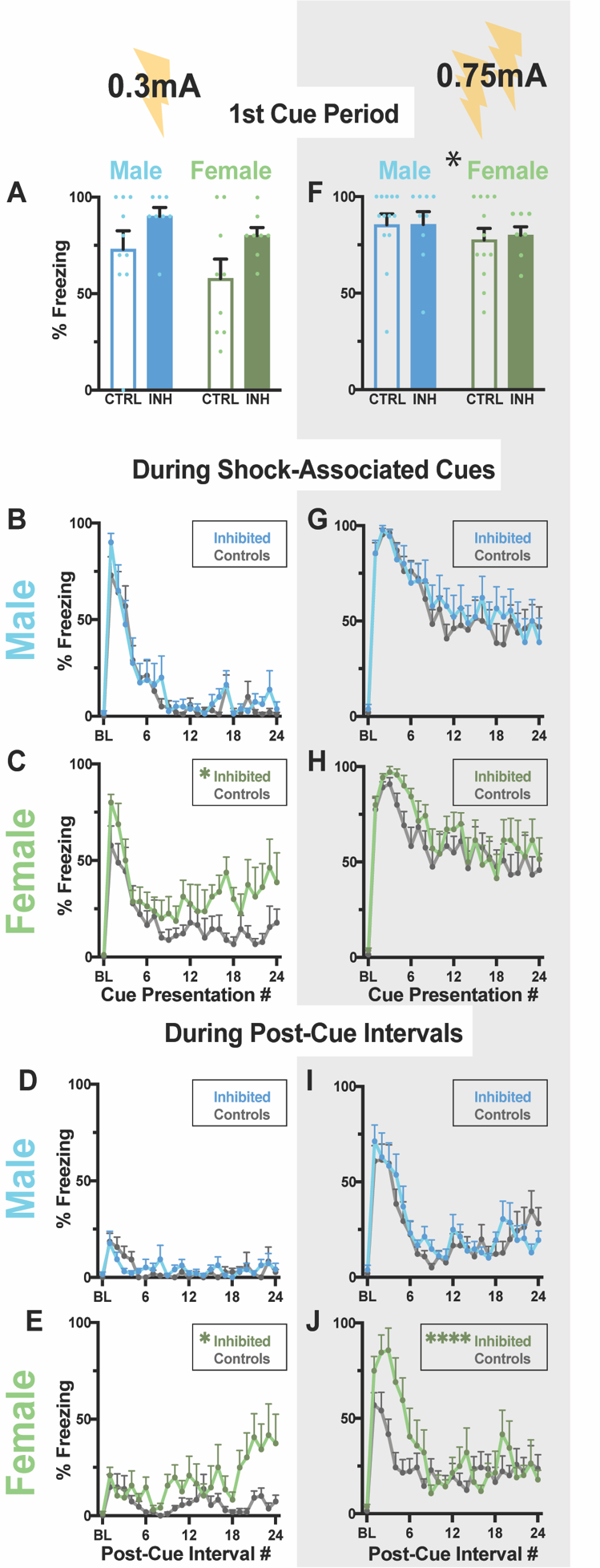

**Supplemental Figure 4: Female Rats Exhibit More Persistent Enhancement of Freezing Following VP^GABA^ Inhibition Than Males.** In the fear expression test for the low-shock cohort, (A) Inhibition enhances freezing to the 1^st^ cue similarly in both sexes, but only females showed enhanced freezing (B, C) during subsequent cue presentations, and (D, E) in post-cue periods. For the high-shock cohort, Inhibition had no effects on (F) 1^st^ cue freezing, or (G,H) freezing during subsequent cues in either sex, but (I, J) only Inhibited females showed significant post-cue enhancement of freezing. INH: Inhibited. CTRL: Controls. Main effect of Sex (Black *) or Inhibition (Green *), p<0.05.

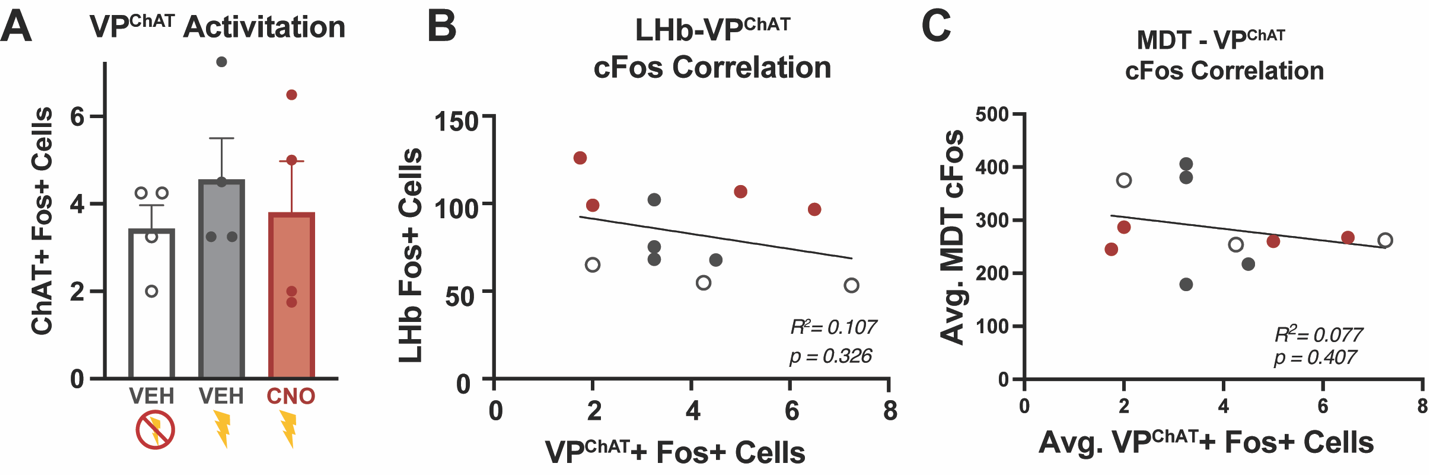

**Supplemental Figure 5: Fos Analysis in VP^ChAT^ cells.** Fos quantification of the No-Threat:VEH, Threat-VEH, and Threat-CNO groups in (A) ChAT-expressing VP cells. VP^ChAT^ Fos does not correlate with Fos in (B) LHb or (C) MDT of the same rats.
